# Myo10-dependent leading-edge coordination maintains physical coherence of migrating dendritic cells

**DOI:** 10.64898/2026.10.17.755813

**Authors:** Anjelika Gasilina, Paniz Rezvan Sangsari, Natalie M. Badillo, Yang-In Yim, Dillon C. Shrock, Nicole Y. Morgan, John A. Hammer

## Abstract

Dendritic cells (DCs) are the principal leukocytes responsible for priming T cells. DC migration through geometrically complex tissue environments is crucial for initiating robust and spatiotemporally coordinated activation of adaptive immune responses. However, our understanding of the molecular mechanisms that govern physical navigation of DCs remains incomplete. Here we identify the unconventional myosin Myo10 as a critical regulator of DC trafficking. We show that loss of Myo10 attenuates FITC-induced contact hypersensitivity and reduces the efficiency of *in vivo* DC migration and egress from dermal explants but notably has no effect on antigen uptake or *in vitro* T cell priming. We further find that the migration defects of Myo10-deficient cells are exacerbated by increasing geometrical complexity. Mechanistically, Myo10 promotes physical coherence and coordinated leading edge dynamics via its interaction with lamellipodin, which subsequently drives VASP-mediated actin polymerization. Our findings identify a Myo10-lamellipodin-VASP axis critical for leukocyte navigation through restrictive tissue architectures, linking cytoskeletal coordination to effect immune surveillance.

## Introduction

Migratory dendritic cells (DCs) act as the primary sentinels of the immune system, whose role is to initiate antigen-specific adaptive immune responses to foreign antigens and maintain tolerance to self-antigens (*1, 2*). This critical role relies on their superior ability to take up antigen, traffic from peripheral tissue to draining lymph nodes, and subsequently prime T lymphocytes. Because interstitial trafficking constitutes the rate-limiting step in the induction of adaptive immunity, it must be executed with precise spatiotemporal coordination. Failures or delays in DC mobilization compromise immune surveillance, leading to unresolved infections, impaired tolerance, and autoimmunity (*3–6*). To accomplish this rapid passage, DCs must successfully navigate three-dimensionally complex, physically confining environments.

Directed migration of DCs to the tissue-draining lymph nodes is guided by CCR7 signaling in response to CCL19 and CCL21 chemokine gradients (*7–9*). Chemokine receptor engagement triggers symmetry breaking and establishes robust front-to-back polarity, driven by RAC- and CDC42-mediated actin polymerization at the cell front and RHOA-dependent non-muscle myosin 2A (NM2A) contractility at the trailing edge. Polarity is stabilized by a localized positive feedback loop wherein spatial accumulation of PtdIns(3,4,5)P_3_ drives localized actin assembly(*10–13*). To traffic in diverse physiological terrains, DCs and other leukocytes switch between integrin adhesion-dependent and integrin adhesion-independent migration modes.

Under physical constraints of complex 3D environments, DCs adopt an integrin adhesion-independent “ameboid” mode of migration, relying on rapid shape changes to push against tissue topology (*14–18*). Ameboid navigation is driven by two spatially segregated forces – localized actin polymerization that drives leading-edge protrusion and NM2A-mediated contractility that pushes the cell rear. Migration in geometrically complex tissues force migrating DCs to break the uniform leading edge into multiple competing protrusions. To prevent cell fragmentation, cells must continuously coordinate protrusion extension, pore size selection, retraction of losing protrusions, reestablishment of a uniform leading edge, and retraction and forward translocation of the cell’s rear. When competing fronts are presented with pores of different sizes, DCs employ the nucleus as a mechanical gauge to select the optimal path. Once the nucleus selects and passes through the largest pore, the corresponding protrusion becomes the winner and all other competing protrusions retract, reestablishing a uniform leading edge (*18–21*). However, when pore dimensions are equivalent, protrusion dominance is determined by higher local accumulation of PtdIns(3,4,5)P_3_, elevated CDC42 activation, and higher local actin polymerization. In both scenarios NM2A-driven contractility at the trailing edge and in the losing protrusions reinforces cell rear retraction and maintenance of polarity (*22, 23*).

Seminal studies from the Halin, Lennon-Dumenil, and Sixt labs elucidated the biochemical and biophysical principles of DC antigen capture and migration through complex environments (*7, 15–18, 20, 21, 24–30*). Relevant to this study, work from the Sixt laboratory demonstrated that cell coherence during pathfinding is regulated by a microtubule-dependent “proprioceptive” feedback loop, wherein local microtubule depolymerization releases RhoGEF Lfc (GEF-H1) to trigger actomyosin-mediated retraction of losing protrusions (*18, 21*). While this process explains how the losing protrusions are dismantled, the complementary molecular machinery that enforces maintenance of the leading edge and prioritizes the winner protrusion remains poorly understood. A central candidate in this process is the small GTPase CDC42, which indispensable for maintaining leukocyte polarity and coordinated migration (*16, 23, 31*).

Intriguingly, foundational studies by Cheney and colleagues established that active CDC42 works alongside the unconventional actin motor Myosin-10 (Myo10, Myosin-X) to drive the initiation of filopodia (*32*). Given that unconventional myosin Myo10 is recruited to and activated at the leading edge *via* its interaction with PtdIns(3,4,5)P_3_, and coordinates actin elongation factors - including Ena/VASP proteins that synergize with the WAVE regulatory complex (WRC) to drive Arp2/3-mediated branched actin assembly - we hypothesized that Myo10 functions as a leading-edge coordinator to preserve DC physical coherence during amoeboid navigation (*32–38*).

Here, we identify Myo10 as an essential regulator of DC ameboid trafficking and adaptive immune activation. Using *in vivo* and *ex vivo* models, we show that genetic deletion of Myo10 attenuates FITC-induced contact hypersensitivity and markedly reduces both DC egress from the dermis and transit to draining lymph nodes, without altering antigen uptake or *in vitro* T cell priming capacity. By challenging DCs with environments of varying physical restriction, we reveal that the migration defect of Myo10-deficient DCs scales directly with increasing geometrical complexity. In the absence of Myo10, DCs fail to maintain directional persistence, losing physical coherence. Mechanistically, we establish the Myo10 maintains leading edge dominance by its interaction with lamellipodin (Lpd), thereby spatially coordinating VASP-mediated actin polymerization at the winning protrusion. Together, our findings define a Myo10-Lpd-VASP axis that coordinates protrusion dynamics to preserve physical coherence during leukocyte navigation through restrictive tissue microenvironments.

## Results

### Myo10 deficiency attenuates cutaneous adaptive immune response due to impaired trafficking of dendritic cells to draining lymph nodes (dLNs)

Mounting an effective adaptive immune response requires a coordinated sequence – capture of peripheral antigen, migration of DCs to the dLNs, presentation of the antigen to naïve T cells and subsequent expansion of antigen-specific effector T cells. To determine whether Myo10 is required for adaptive immune activation in vivo, we evaluated wild-type (WT) and Myo10-deficient (KO) mice using a fluorescein isothiocyanate (FITC)-induced contact hypersensitivity (CHS) model, a well-established assay of cutaneous Th2-skewed inflammation. Mice were sensitized on the flanks with topical FITC (0.5% in dibutyl phthalate/acetone) on days 0 and 6 and challenged on the dorsal side of one ear on days 12, 13, and 14 (Fig. 1A). Thickness of challenged and unchallenged ears was measured to monitor the onset and magnitude of local inflammation. Whereas WT mice exhibited progressive increase in ear swelling from 0.203 ± 0.016 mm on day 12 to 0.562 ± 0.035 mm on day 15, Myo10 KO mice displayed an attenuated response, reaching only 0.400 ± 0.013 mm on day 15 (Fig. 1B). These findings indicate that loss of Myo10 results in an attenuated FITC-induced cutaneous immune response.

**Figure 1.**
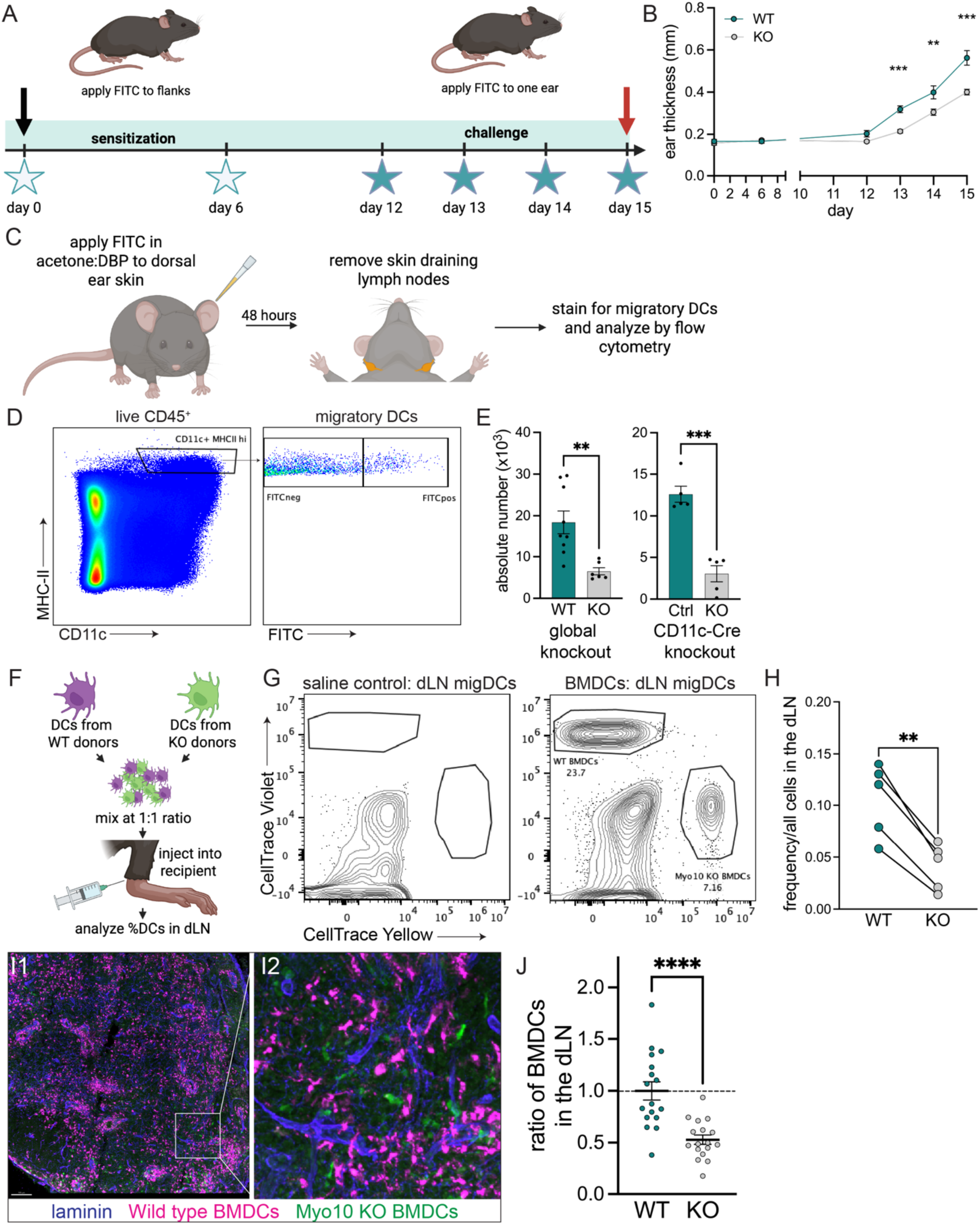
Myo10 depletion attenuates adaptive immune response due to impaired migration of dendritic cells to the skin dLNs. **(A - B)** FITC-induced contact hypersensitivity response (CHS) in wild type (WT) and global Myo10 knockout (KO) mice. **(A)** Schematic depiction of the experiment. **(B)** Graph summarizing quantitation of ear thickness (mm) measured on the indicated days in WT and Myo10 KO mice. Data are plotted as mean ± SE (** p< 0.01, ***p<0.0001, based on two-way ANOVA with multiple comparisons using Šídák’s correction). n = 5 mice per group. **(C - E)** Results of FITC painting experiments. Ears of wild type or Myo10^-/-^ mice were topically treated with 1% FITC/dibutyl phthalate:acetone mixture and dLNs were analyzed by flow cytometry. **(C)** Schematic depiction of the experiment. **(D)** Flow cytometry gating scheme on auricular dLNs. **(E)** Graphs summarizing quantitation of absolute numbers of FITC-positive migratory (CD11c^+^ MHCII^hi^) DCs from global and CD11c-Cre-driven knockout models. Each dot represents a value from one mouse. Data are plotted as mean ± SE, ** p< 0.01, ***p<0.0001 based on unpaired t-test. **(F – J)** Results of adoptive transfer experiments. BMDCs arriving in the draining lymph node (dLN) 48 hours later were analyzed by flow cytometry **(G – H)** or confocal microscopy (I – J). (F) Schematic depiction of the experiment. **(G)** Flow cytometry plot showing gating and resulting percentages of WT and KO BMDCs in the popliteal dLN suspension. **(H)** Graph summarizing quantitation of frequencies of adoptively transferred BMDCs. Each dot represents a value from one recipient mouse. Data are plotted as mean ± SE (** p< 0.01 based on a paired t-test). Each dot represents a value from one mouse, n = 5 mice. **(I1)** Confocal image of a cryosection of a laminin-counterstained (blue) dLN showing arrival of wild type (magenta) and Myo10 KO (green) BMDCs and **(I2)** enlarged area of the inset shown in white region of interest in **I1**. scale bar = 100 µm. **(J)** Quantitation of BMDCs present in the lymph node relative to wild type. n = 9 recipient mice. Data are plotted as mean ± SE, **** p< 0.0001 based on an unpaired t-test.

Because productive adaptive immune response requires efficient DC transit from the periphery to dLNs, we next asked whether this observed blunted CHS response arose from defective DC trafficking. We performed a FITC painting assay by applying topical 0.5% FITC to the dorsal sides of ears of WT and KO mice and quantitating the absolute numbers of migrated DCs in auricular dLNs. (Fig. 1C). Flow cytometric analysis of migratory DCs (CD11c^+^, MHC-II^hi^) revealed a marked reduction in FITC^+^ DCs arriving in the dLNs of KO mice compared to WT controls (6.50 x 10^6^ ± 0.86 x 10^6^ KO *vs.* 18.42 x 10^6^ ± 2.74 x 10^6^ WT; Fig. 1D, see also Fig. S1B for gating strategy). To determine whether this phenotype is restricted to CD11c lineage, we generated a CD11c-specific Myo10 KO. In agreement with the results in the global KO, CD11c-specific Myo10 KO model displayed fewer migratory DCs in skin dLNs than the littermate controls (3.04 x 10^6^ ± 0.97 x 10^6^ CD11C-Cre-KO vs. 12.6 x 10^6^ ± 0.96 x 10^6^ littermate control) (Fig. 1E). These results demonstrate that Myo10 is critical for migration of DCs in response to cutaneous challenge.

To rule out potential contributions from altered stromal microenvironments in KO mice, we performed a competitive *in vivo* homing assay. Fluorescently labeled WT and KO bone marrow derived DCs (BMDCs) were co-injected at 1:1 ratio into the lateral tarsal region of the hind foot (Fig. 1F). Flow cytometric analysis of popliteal dLNs 48 hours post-transfer revealed a significant underrepresentation of KO BMDCs among migratory DC (CD11c^+^, MHC-II^hi^) cells compared to the WT (8.19 ± 1.08% for KO *vs.* 26.30 ± 0.74% for WT); (Fig. 1G-H, S1A).

Confocal analysis of cryosections of popliteal dLNs further confirmed reduced ratio of Myo10 KO:WT DCs (Fig. 1I, J) in the lymph node. These results demonstrate that Myo10 is critical for efficient homing of DCs. Considering that our *in vitro* data showed no appreciable differences in antigen uptake between WT and Myo10 KO DCs (Fig. S1C-G) and no appreciable differences in the surface expression of CD86, CD80, and MHC-II (Fig. S1H, I), the results of our *in vivo* experiments demonstrate that Myo10 selectively regulates directed migration of DCs.

### Myo10 is necessary for interstitial migration, and lymphatic vessel entry of migrating DCs and their directional persistence in fibrous matrices

Interstitial DC trafficking from peripheral skin to draining lymph nodes requires cells to navigate connective tissue networks of varying density and geometry before entering lymphatic vessels. To isolate where DC trafficking fails in the absence of Myo10, we used *ex vivo* crawl-out assays (*39, 40*), which quantitate the emigration of endogenous skin-resident DC subsets from dermal tissue into culture medium (Fig. 2A). Through flow cytometric quantitation of absolute numbers of migratory DCs present in the media and retained in the ear after 48 hours (Fig. 2B and S2A) we observed a significantly smaller ratio of endogenous DC, in which Myo10 was knocked out, in the media compared to the respective control animal ear explants (0.37 ± 0.11 *vs.* 0.73 ± 0.11 for CD11c-lineage KO *vs.* littermate controls and 0.54 ± 0.02 *vs.* 0.83 ± 0.02 for global KO *vs.* WT) (Fig. 2 C-D). These results suggest that without Myo10, DCs fail to efficiently emigrate from the dermis.

**Figure 2.**
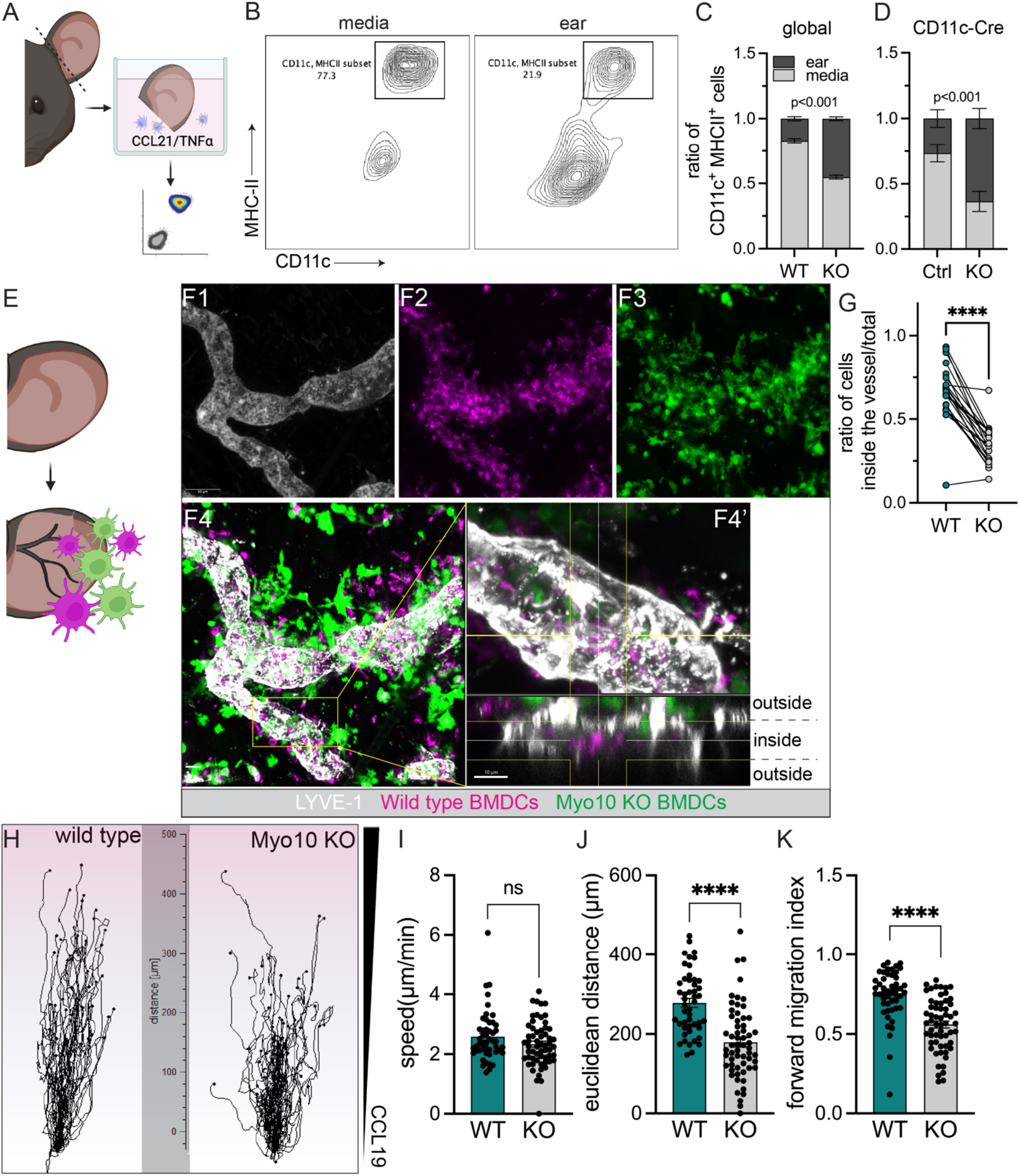
Myo10 loss impedes dendritic cell egress from the dermis. **(A - D)** Results of the endogenous ear crawl-out assays. Ears were split along the cartilage and floated dermis side down in media containing CCL21 and TNFα. Egress of skin endogenous DC populations was quantitated using flow cytometric analysis 48 hours later. **(A)** Schematic depiction of the experiment. **(B)** Flow cytometry plots depict the gating scheme of emigrated (media) and retained (ear) DCs. Absolute numbers of DCs in the media and in the ear 48 hours post-floatation in **(C)** WT and global Myo10 KO model and **(D)** Control and CD11c-Cre-driven Myo10 KO model. **(E - G)** Results of exogenous ear crawl-in assays. **(E)** Schematic summarizing the experimental setup. CellTrace e670-labeled WT BMDCs and CellTracker Green-labeled Myo10 KO BMDCs were mixed at 1:1 ratio and placed on top of the exposed dermis. Three hours post incubation rinsed ear halves were fixed and processed for confocal microscopy. **(F)** Representative whole-mount images showing localization of wild type (magenta) and Myo10 KO (green) BMDCs. Single channel images show **(F1)** lymphatic vessel labeled with LYVE-1 antibody, **(F2)** localization of WT BMDCs, **(F3)** localization of KO BMDCs. **(F4 and F4’)** are channel-merged images showing localization of WT and KO BMDCs relative to the lumen. Scale bar = 20 µm **(G)** Graph summarizing quantitation of ratios of BMDCs situated within the vessel, n = 6 recipient mice and 4 independent BMDC cultures. Data are plotted as means ± SE, ****p<0.0001 based on a paired t-test. **(H - K)** Results of an in vitro 3D collagen migration assay. **(H)** Spider plot showing single cell tracks of migrating BMDCs. Graphs summarizing quantitation of speeds **(I)**, Euclidean distance **(J)**, and forward migration index **(K)** of chemotaxing BMDCs. Each dot represents a value from one cell. Data are combined from 3 independent experiments and plotted as mean ± SE, n = 10 – 30 cells per condition per experiment

To dissect whether this defect arises during interstitial matrix transit or/and lymphatic vessel entry, we performed a competitive dermal crawl-in assay. Differentially labeled WT and KO BMDCs were mixed in an equimolar ratio and seeded directly on top of the exposed dermis of a dorsal ear half of murine ear (Fig. 2E). Following a 30 min incubation, the ear halves were rinsed to remove any cells that didn’t enter the dermis and further incubated for up to 3 hours in complete media. Confocal microscopy quantitation of BMDCs within the vessel lumen (marked by LYVE-1) (Fig. 2F1-F4; Fig. S2 B1-B3) revealed lower ratio of KO BMDCs compared to the WT (0.34 ± 0.02 *vs.* 0.66 ± 0.03, respectively) (Fig. 2G). Notably, we observed that many KO BMDCs were present in the collagen-rich dermis surrounding the lymphatic vessels.

We posited that DCs lacking Myo10 are unable to migrate within dense fibrillar collagen networks. We performed migration assays in collagen gels using murine tissue derived primary DCs (Fig. 2H-K; S2 C1-C3) and BMDCs (Fig. S3 B-E) from WT and Myo10 KO animals. Although we observed no appreciable differences in speeds between the WT and Myo10 KO DCs (2.56 ± 0.12 µm/min *vs.* 2.34 ± 0.10 µm/min), we saw significant reduction in mean Euclidian distance of 277.9 ± 11.32 µm *vs.* 179.1 ± 11.76 µm and forward migration index (0.75 ± 0.02 vs. 0.56 ± 0.02) (Fig. 2I-K). We observed similar phenomena in invasion and migration though collagen gels in BMDCs. Quantitation of the endpoint analysis of cells invading into the collagen I gel showed that WT BMDCs traveled a mean distance of 805.7 ± 28.04 µm compared to Myo10 KO BMDCs, which migrated a mean distance of 579.2 ± 25.45 µm (Fig. S2F-G). Flow cytometric evaluation of CCR7 expression revealed no meaningful differences, suggesting no defect in chemokine sensing (Fig. S3 H-I). These results suggest that while Myo10 is dispensable for motility and the initiation of chemotaxis, it is necessary for the maintenance of directional persistence and efficient interstitial invasion within fibrous 3D microenvironments.

### Migration defects of Myo10 KO DCs scale with geometric complexity and pore size heterogeneity

The ability of migrating DCs, and leukocytes in general, to efficiently navigate tissues requires continuous coordination between cell polarity, directional persistence, and response to physical constraints imposed by the surrounding environment. Because dermal explants and *in vitro* collagen matrices provide only modest control over pore sizes and geometries, we developed and fabricated microfluidic channels with defined dimensions and branching architecture to systematically vary confinement, topology, and branching architecture.

We first built bifurcated microchannels containing serially repeating junctions (Fig. 3A). This design required that individual BMDCs repeatedly encounter a choice point, project competing protrusions into alternative paths, select a direction, retract the non-productive protrusion, and subsequently repeat this process at each junction to sustain forward translocation. Both WT and KO BMDCs entered the channels readily (Fig. 3B), however, KO BMDCs exhibited an increased passage time of 52.24 ± 5.59 min *vs.* WT of 27.05 ± 1.98 min (Fig. 3C). Moreover, KO BMDCs displayed a significantly increased aspect ratio compared to WT BMDCs (average length of 128.4 ± 3.58 µm for WT *vs.* 192.4 ± 9.99 µM for KO) (Fig. 3D). To further assess the ability of BMDCs to negotiate repeated junctions, we quantitated the number of junctions traversed by each cell during the imaging period. Only 3.34 ± 1.14% of KO BMDCs successfully passed through 5 or more obstacles, compared to 40.82 ± 7.11% WT BMDCs (Fig. 3E).

**Figure 3.**
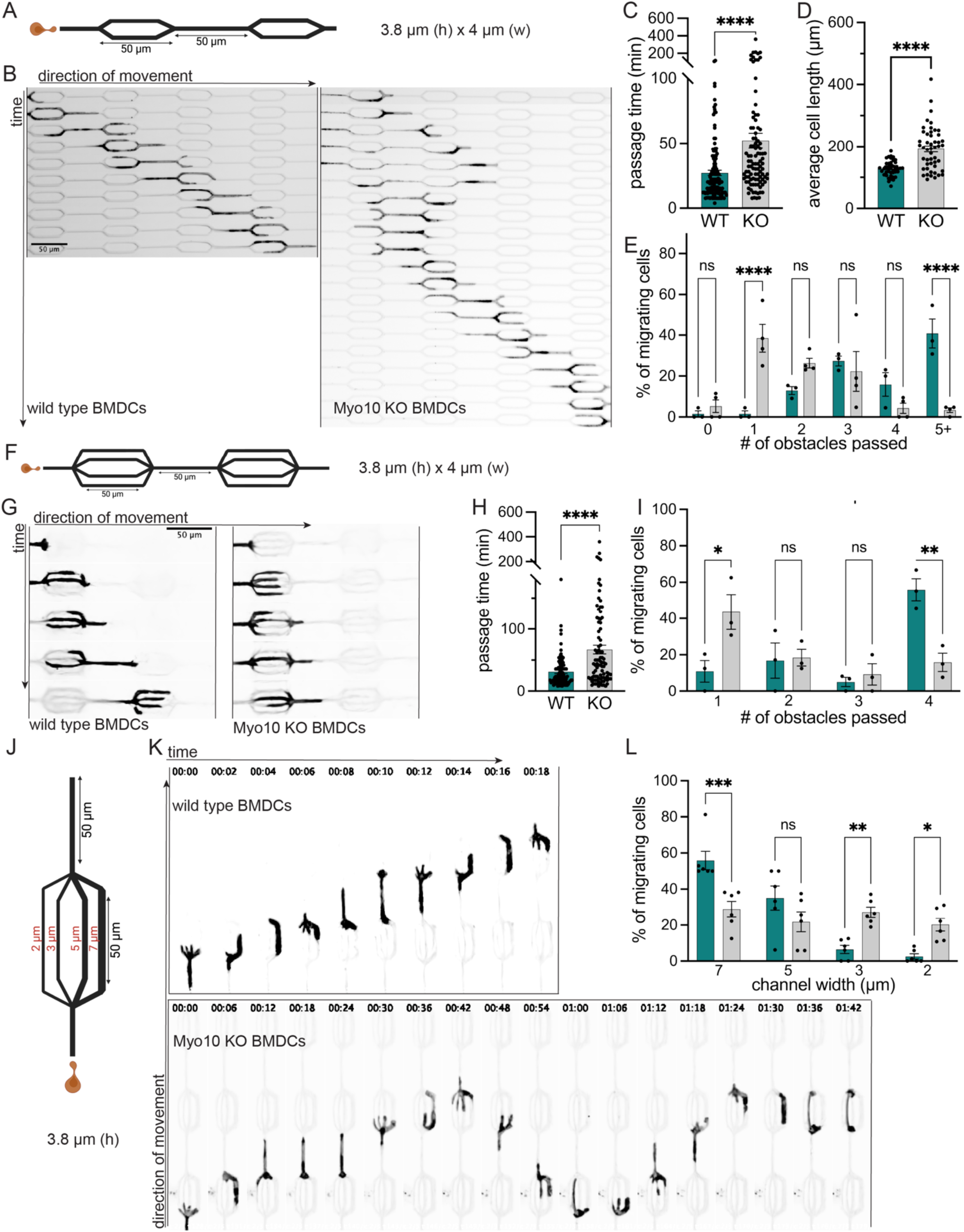
Myo10 deficiency impairs DC migratory persistence and decision-making in complex environments. **(A – E)** Migration dynamics and efficiency in 2-way (bifurcated) microfabricated channels. **(A)** Schematic representation of the bifurcated microchannel design with serially repeating 2-way junctions. **(B)** Montage of a migrating WT and KO cell from a representative experiment. **(C)** Graph summarizing quantitation of the channel passage time (in min) for WT and KO BMDCs. Data are presented as means ± SE (****p<0.0001 based on two-tailed unpaired t-test). n = 130 – 140 cells per genotype pooled from 3 independent experiments. **(D)** Graph summarizing quantitation of the cells’ aspect ratio during decision-making and passage. Data are presented as means ± SE (****p<0.0001 based on two-tailed unpaired t-test). n = 40 - 45 cells per genotype pooled from 3 independent experiments. **(E)** Graph summarizing quantitation of percentage of migrating cells traversing one or more continuous junction in an imaging period. Data are plotted as means ± SE (****p<0.0001 based on two-way ANOVA with multiple comparisons using Šídák’s correction. **(F – I)** Migration dynamics and efficiency in 4 – way (quadfurcated) microfabricated channels. **(F)** Schematic representation of the quadfurcated microchannel design, showing confinement height, width, and junction lengths**. (G)** Montage of a migrating WT and KO cell from a representative experiment. **(H)** Graph summarizing quantitation of junction passage time (in min) per 4 – way obstacle for WT and KO BMDCs. **(I)** Graph summarizing percentage of WT and KO BMDCs successfully traversing junctions. Data are presented as means ± SE (ns – not significant, **p>0.01, ****p<0.0001 based on two-way ANOVA with multiple comparisons using Šídák’s correction). n = 3 independent experiments. **(J - L)** Migration dynamics and efficiency in 4 – way (quadfurcated) microfabricated channels with varying fork widths. **(J)** Schematic representation of the channel design. **(K)** Montage of a migrating WT and KO cell from a representative experiment. **(L)** Pore size preference of migrating BMDCs

We posited that the migratory defect will become more pronounced with increased complexity of the environment, therefore we developed and built microchannels containing 4-way repeating junctions (Fig. 3F). Here, the quadfurcated design also required the cells to repeatedly polarize, extend protrusions, select a path, retract non-productive protrusions, however, the increased number of forks at each junction placed a greater weight on decision - making and leading edge coordination. Unlike WT BMDCs, which passed through each junction with an average passage time of 31.35 ± 2.63 min, KO BMDCs required 66.90 ± 6.99 min to pass each junction (Fig. 3G-H). Due to prolonged entrapment at junctions caused by multiple competing leading edges, only 15.90 ± 5.02% of KO BMDCs successfully traversed 4 or more obstacles, compared to 55.89 ± 6.03% WT BMDCs (Fig. 3I).

Renkawitz et al. demonstrated that leukocytes, such as BMDCs, navigate complex environments by preferentially using the path of least resistance. Therefore, we asked whether KO BMDCs retain the ability to select more physically permissive migration path by building 4-way microchannels, in which each junction presents four forks, each of defined – 2, 3, 5, or 7 µm – width (Fig. 3J). In agreement with previously published observations (*18*), WT DCs showed a strong pore size preference (Fig. 3K), with majority of cells selecting a fork with the largest width - 55.90 ± 5.1% cells and 35.07 ± 6.7% of cells selecting channel size 7 µm and 5 µm, respectively (Fig. 3L). In contrast, KO cells showed no significant pore size preference, with cells choosing indiscriminately – 20.25 ± 3.57% committing to a 2 µm fork, 21.18 ± 2.80% committing to a 3 µm, 21.83 ± 5.52% committing to a 5 µm, and 28.90 ± 4.3% committing to a 7 µm fork. These results suggest that unlike WT DCs, KO DCs no longer preferentially chose the path of least resistance. Collectively, our data in microfabricated devices suggest that Myo10 KO BMDCs can generate exploratory protrusions and initiate forward movement but exhibit impaired directional persistence, decision making, and selection of permissive migratory paths, all of which results in inefficient navigation through confined environments.

### Myo10 maintains leading edge coherence and restricts protrusive entropy during confined migration

Having established that Myo10 is needed for directional navigation, we next examined the leading edge dynamics directly to determine whether the impaired directional migration of KO DCs stemmed from altered organization of the leading edge. We turned to under agarose assays, which provide uniform vertical confinement. Consistent with the phenotype observed in collagen gels and microchannels, KO BMDCs frequently displayed unstable front-rear polarity and formation of multiple competing leading edges. Whereas WT cells typically maintained a single dominant leading edge that supported continuous forward progression (Fig. 4A1), KO cells frequently generated multiple competing protrusions that persisted or shifted abruptly in orientation during migration (Fig. 4A2-3; S4A and C1-C2). Quantitative analysis of trajectories and migratory parameters revealed less coherent migration by KO cells (Fig. 4B-E; S4B and S4D-G).

**Figure 4.**
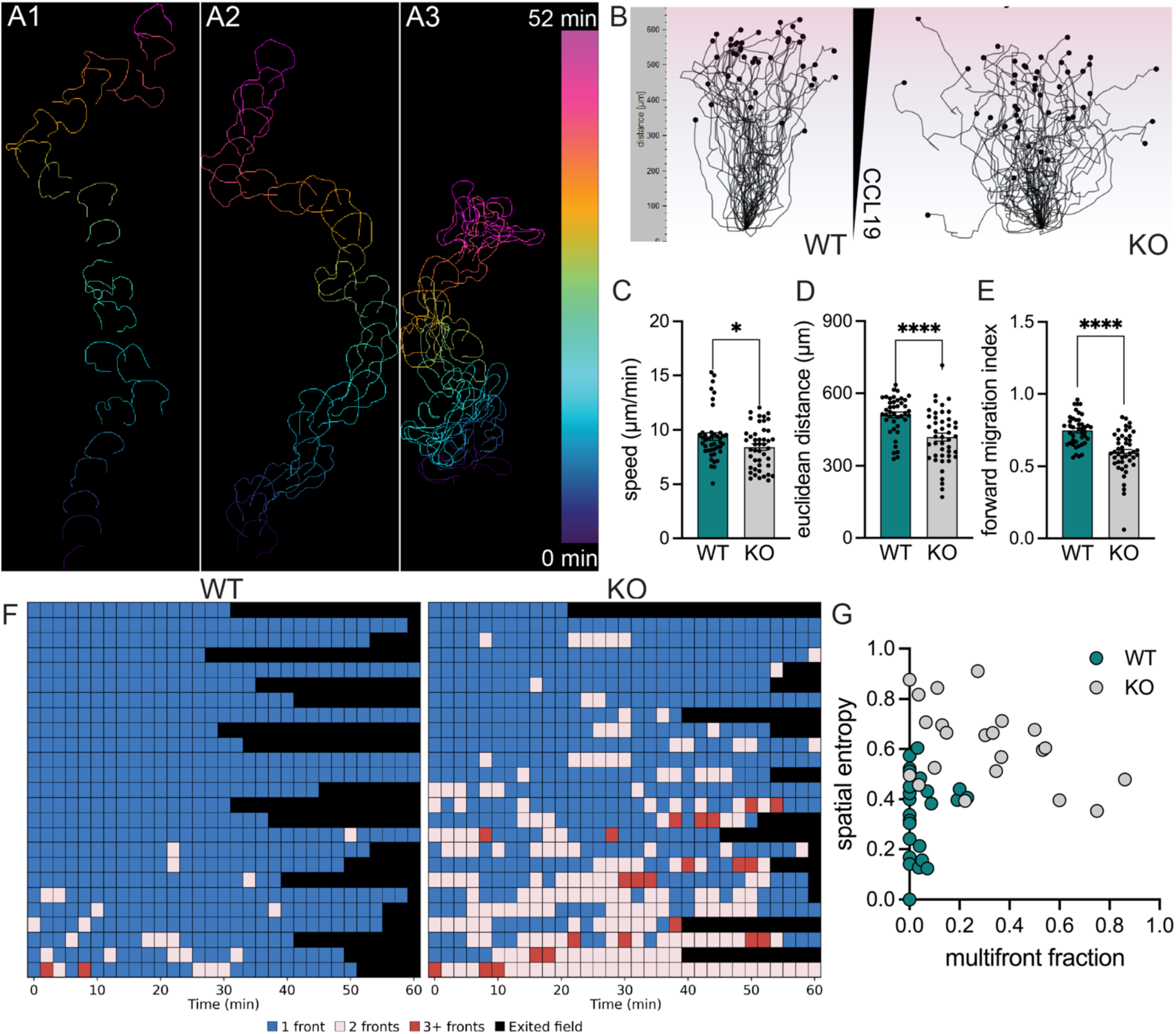
Myo10 maintains for leading edge stability and directionality. **(A1-A3)** Temporal projection of representative WT **(A1)** and Myo10-deficient (KO) (**A2** – example of productive migration, **A2** – example of non-productive migration) BMDCs migrating under agarose toward CCL19. **(B)** Origin-normalized migration trajectories of WT and Myo10 KO BMDCs. Dots indicate the terminal position of individual cell tracks. **(C–E)** Quantitation of migration parameters in WT and Myo10 KO BMDCs, including **(C)** speed, **(D)** Euclidian distance, and **(E)** forward migration index. Each dot represents an individual cell. **(F)** Temporal analysis of leading-edge organization in individual WT and Myo10 KO cells. Each row represents one cell and each column represents a time point. Leading edge states were classified as one front (blue), two fronts (pink), or three or more fronts (red). Black indicates time points after a cell exited the field of view and could no longer be tracked. **(G)** Graph summarizing relationship between multi-front occupancy and spatial entropy of leading edge activity for individual WT and KO cells. Multifront fraction = proportion of analyzed time points containing two or more simultaneous fronts. Spatial entropy = normalized Shannon entropy of dominant front occupancy, with increasing values indicating greater spatial redistribution of the dominant front. Each dot represents an individual cell.

To define the temporal stability of the leading edge, we quantitated protrusion architecture across individual migration tracks by classifying each time point according to the number of active, simultaneously advancing fronts. State transition analysis over time revealed that WT cells predominantly occupied a single-front state, whereas KO cells repeatedly occupied two- or multi-front states and transitioned between leading edge configurations (Fig. 4F). Thus, loss of Myo10 produces altered the temporal stability of leading edge organization.

To distinguish the frequency of competing protrusions from the overall spatiotemporal disorder of front activity, we plotted the fraction of multi-front observations against the spatial entropy of leading edge protrusion for individual cells. WT cells clustered within a low multi-front, low entropy region, indicative of a spatially organized, stable leading edge. Conversely, KO cells exhibited a pronounced shift toward high multi-front occupancy accompanied by elevated spatial entropy (Fig. 4G). These results indicate that Myo10 coordinates protrusive activity, restricts spatial entropy, and enforces leading edge coherence.

### Myo10 couples polarized PtdIns(3,4,5)P_3_ signaling to actin assembly at the leading edge

Persistent cell migration requires the establishment and maintenance of a spatially restricted leading edge, where signaling and actin assembly are coupled to sustain protrusion.

Because Myo10 KO DCs frequently maintained competing protrusions, we asked whether actin polymerization was impaired within these multiple fronts (Fig. 5A-B). Quantitative analysis of fluorescent phalloidin revealed significantly reduced F-actin content at the immediate boundary of the leading edge of KO cells compared with WT cells (Fig. 5C).

**Figure 5.**
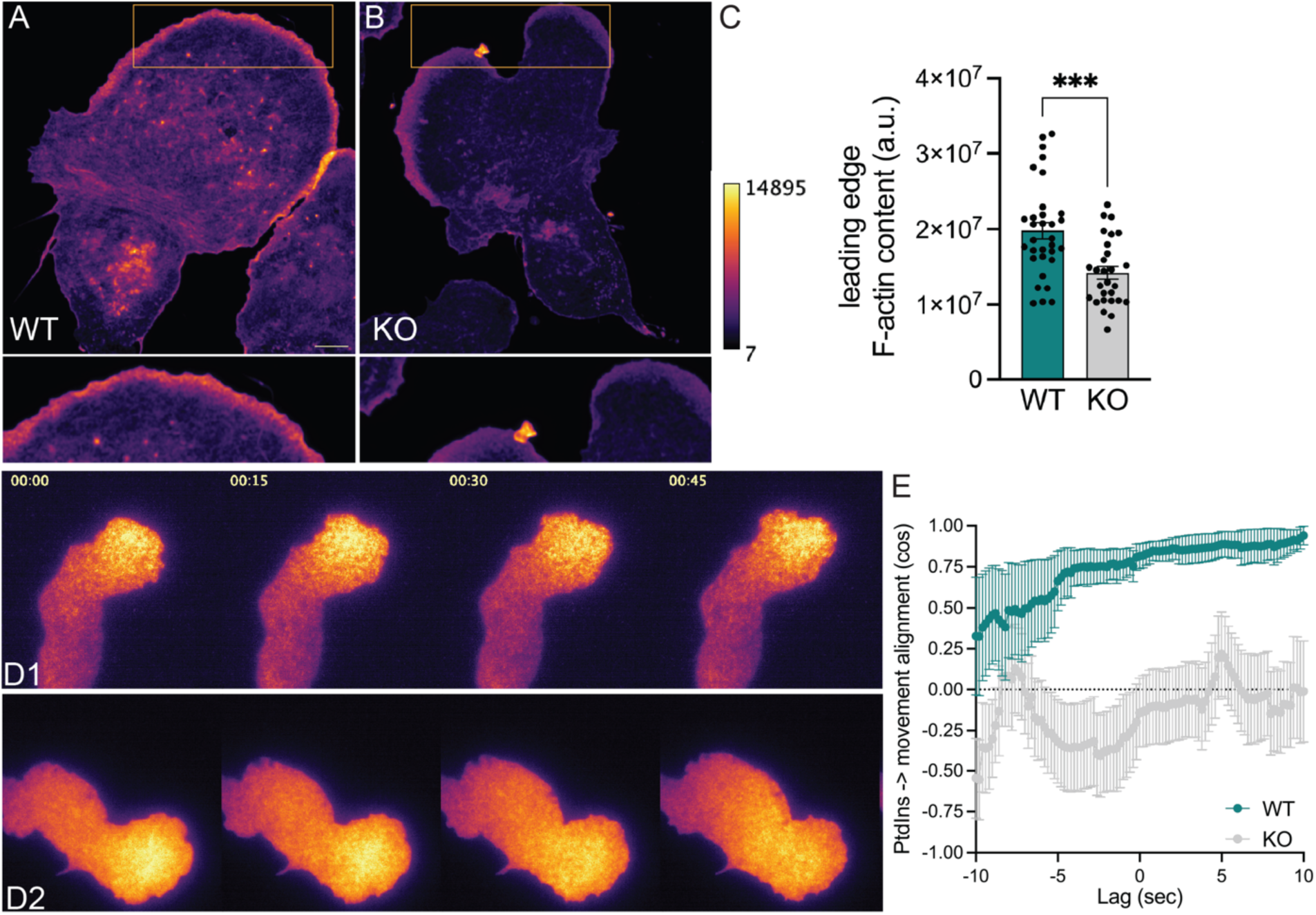
Myo10 coordinates leading edge F-actin assembly with polarized PtdIns(3,4,5)P_3_ signaling and directional movement. **(A - B)** Representative phalloidin stained WT **(A)** and KO **(B)** BMDCs showing F-actin distribution. Yellow boxes indicate the leading edge regions shown at higher magnification below. Images are displayed using the indicated fluorescence intensity scale. **(C)** Quantitation of F-actin content within the leading-edge region of WT and Myo10 KO BMDCs. Each point represents an individual cell. **(D)** Snapshots of representative time-lapse videos of PtdIns(3,4,5)P_3_ dynamics in migrating WT **(D1)** and KO **(D2)** BMDCs expressing AKT-PH-mStayGold. Time stamp = min:sec. **(E)** Temporal alignment between PIP3 polarity and subsequent cell movement in WT and KO BMDCs. Values approaching 1 indicate movement aligned with PIP3 polarity, values near 0 indicate no directional relationship, and negative values indicate movement opposite to PIP3 polarity.

We next determined whether these competing fronts remain coupled to leading edge polarity by evaluating the spatial alignment between the vector of PtdIns(3,4,5)P_3_ enrichment and the direction of subsequent cell translocation (Fig. 5D). In WT DCs, the orientation of peak accumulation was strongly aligned with the vector of cell displacement, with positive cross-correlation increasing across positive temporal lags (Fig. 5E). In contrast, KO cells failed to maintain a predictive relationship between PtdIns(3,4,5)P_3_ orientation and directional movement. Together these results indicate that Myo10 coordinates actin assembly at the leading edge and preserves spatiotemporal coupling between upstream PtdIns(3,4,5)P_3_ signaling and actin-based locomotion.

### Myo10-mediated cell coherence and leading edge coordination are maintained through its interaction with Lpd

Previous work established that Myo10 promotes clustering and accumulation of Ena/VASP proteins at the leading edge and this process is mediated by *via* a direct interaction with Lpd through Myo10C-terminal MyTH4/FERM domain. In parallel, Ena/VASP proteins have been shown to coordinate with the WAVE regulatory complex to enhance Arp2/3-dependent actin assembly (*35, 38, 41–50*). We thus hypothesized that Myo10 functions as a spatial anchor to scaffold Lpd-VASP at the leading edge. We reasoned that depletion of Lpd should phenocopy Myo10 knockout. Indeed, when we knocked down Lpd via Given the defects in leading-edge actin organization and coupling of polarized signaling to directional movement following Myo10 loss, we asked whether Myo10 might regulate protrusive behavior through the actin regulatory protein Lamellipodin (Lpd). Lpd was strongly enriched at the leading edge of migrating control DCs, with fluorescence intensity decreasing from the cell front toward the midbody (Fig. 6A-B). In Myo10-deficient cells, this polarized enrichment was markedly reduced despite preservation of total cellular Lpd protein (Fig. 6C), suggesting that Myo10 contributes to the spatial organization of Lpd rather than its expression.

**Figure 6.**
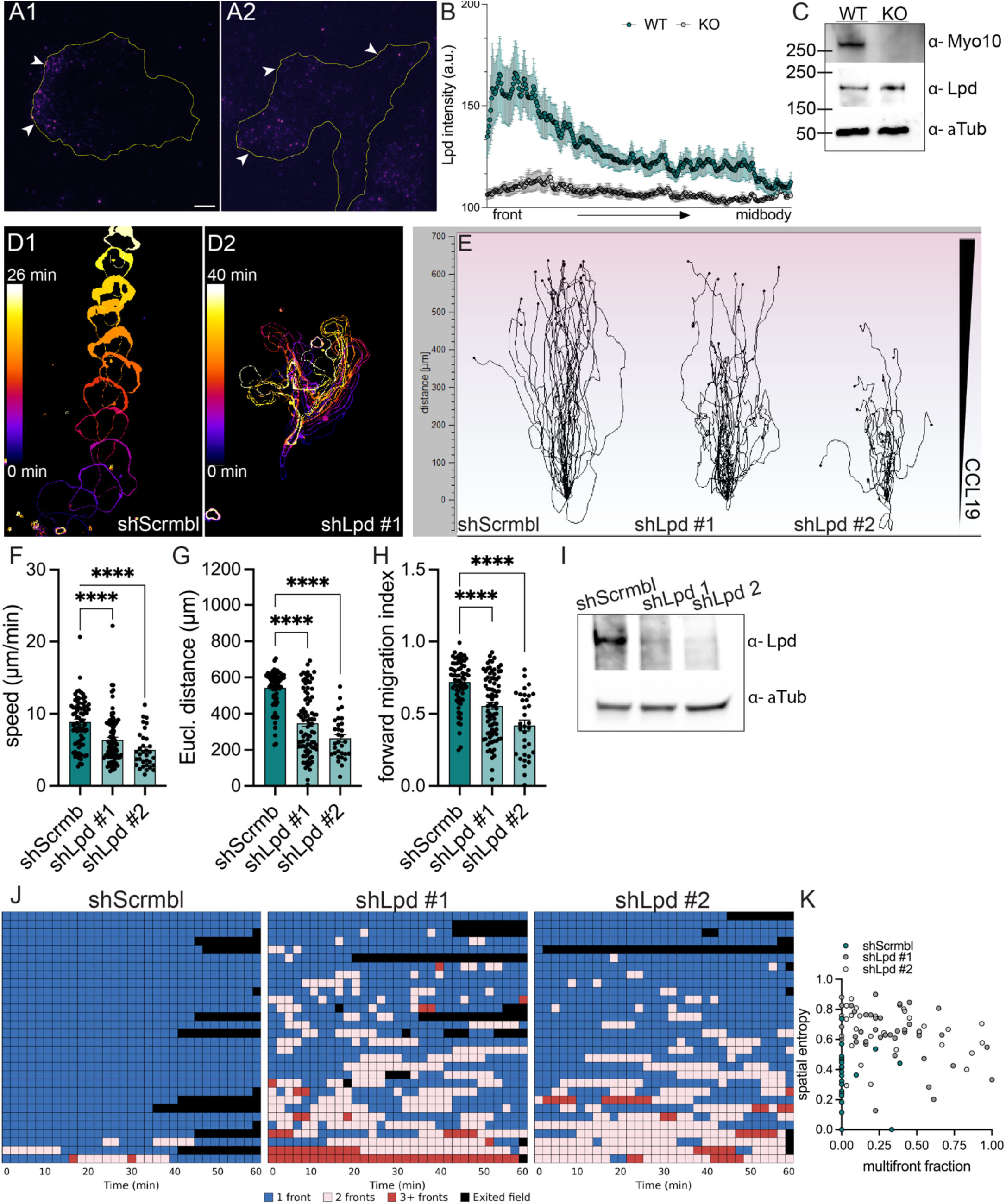
Myo10 - mediated cell coherence and leading edge coordination are maintained through its interaction with Lpd. **(A - C)** Loss of Myo10 blunts recruitment of Lpd to the leading edge of migrating DCs. **(A)** Representative images of chemotaxing WT and Myo10 KO BMDCs (under agarose) stained with Lpd antibody and imaged in TIRF to show clustering of Lpd at the leading edge of WT but not KO cells. **(B)** Graph illustrating quantitation of intensity of Lpd localization in WT and KO leading edges. **(C)** Immunoblot from a representative experiment confirming comparable levels of expression of total Lpd in WT and KO BMDCs. **(D – I)** Downregulation of Lpd phenocopies Myo10 loss. **(D)** Color - coded temporal projections of migrating BMDCs transduced with non-targeting shRNA (D1, shScrbml) or shRNA against mouse Lpd (D2, shLpd #1). **(E)** Single cell tracks of migrating BMDCs stably expressing non-targeting shRNA or shRNA against mouse Lpd. Graphs summarizing quantitation of speeds **(F)**, Eucledian distance **(G)**, and forward migration index **(H)** of chemotaxing BMDCs. Each dot represents a value from one cell. Data are combined from 3 independent experiments and plotted as mean ± SE, n = 10 – 30 cells per condition per experiment. ****p < 0.0001 based on one-way ANOVA with multiple comparisons using Dunnett’s correction. **(I)** Immunoblot from a representative experiment showing knockdown efficiency. **(J-K)** Temporal analysis of leading-edge organization in individual control and shLpd cells. **(J)** Each row represents one cell and each column represents a time point. Leading edge states were classified as one front (blue), two fronts (pink), or three or more fronts (red). Black indicates time points after a cell exited the field of view and could no longer be tracked. **(K)** Graph summarizing relationship between multi-front occupancy and spatial entropy of leading edge activity for individual shScrambl and shLpd cells. Each dot represents an individual cell.

To determine whether Lpd contributes functionally to DC migration, we depleted Lpd using two independent shRNAs (Fig. 6I). Lpd depletion disrupted directional migration toward CCL19, producing disorganized trajectories and significant reductions in migration speed, Euclidean displacement, and forward migration index compared with scrambled-control cells (Fig. 6D–H). We next asked whether Lpd depletion reproduced the leading-edge instability observed following Myo10 loss. Scrambled-control cells predominantly maintained a single leading edge over time, whereas cells depleted of Lpd frequently generated and maintained multiple competing fronts (Fig. 6J). Quantification using the same leading-edge framework applied to Myo10-deficient cells revealed increased multifront behavior together with elevated spatial entropy following Lpd depletion (Fig. 6K). Thus, Lpd depletion phenocopied the loss of leading-edge coherence caused by Myo10 deficiency.

Together, these findings support a model in which Myo10 promotes polarized recruitment or retention of Lpd at the leading edge, enabling protrusive activity to be consolidated into a persistent front and thereby supporting directional migration.

## Discussion

Rapid trafficking of dendritic cells from peripheral tissue to the draining lymph nodes constitutes the rate-limiting step in mounting adaptive immune response. Here we identify the unconventional actin motor protein Myo10 as a critical regulator of this migratory cascade. We show that the loss of Myo10 dampens contact hypersensitivity, reduces accumulation of DCs within dLNs, and impairs egress from tissue without perturbing antigen uptake, uptake-associated maturation, and T cell priming. At the cellular level, these defects were not associated with inability to produce forward locomotion, but rather from a failure to organize migratory activity into a coherent and directionally persistent leading edge. In the absence of Myo10, migrating DCs generate competing fronts with elevated spatial entropy, have decoupled polarized PtdIns(3,4,5)P_3_ - forward locomotion loop. Together our findings define Myo10 as a regulator of productive migratory coherence.

The phenotypic manifestation of Myo10 deficiency scales directly with environmental complexity, highlighting its role in topological decision making. DCs migrating through simple environments can be constrained by the geometry of their surroundings, while geometric junctions and heterogeneous fibrous collagen networks present multiple paths and require cells to project multiple exploratory protrusions and resolve competition to sustain forward movement (*20, 22, 23*). Consistent with models established by Theriot and colleagues (*23, 24, 52*) directional migration can involve transient competition between leading edges, followed by stabilization of the winning front and retraction of the losing front. CDC42 activity can provide a localized polarity signal that precedes directional turning, whereas NM2A activity contributes to remodeling and retraction of the losing front. In our microchannels WT efficiently negotiate multiple branch points, while Myo10 KO cells exhibit prolonged dwell times, hyper-elongated morphologies, and indiscriminate pore size selection. Thus, Myo10 may be particularly important when environmental geometry creates competing states that must be resolved into a single dominant front.

How then does Myo10 function to promote protrusion coordination as DCs migrate through complex environments? In general, unconventional myosin function is dictated by tail domain interactions. For Myo10, this could mean interaction with integrins (*via* its FERM domain), microtubules (*via* its MyTH4 domain), PtdIns(3,4,5)P_3_ (*via* its PH domains), and/or Lamellipodin (Lpd) (*via* a distinct region within the FERM domain). We would argue for the following reasons that Myo10’s interactions with PIP3 and Lamellipodin are interactions that underly Myo10’s role in protrusion coordination. First, integrin binding role is unlikely, because our in vitro assays were conducted on BSA- or PLL-PEG-passivated surfaces, where ameboid motility of DCs is independent of integrin adhesions. Second, microtubule binding seems unlikely to be playing a significant role given the enormous difference in the abundance of Myo10 (very low) and microtubule (very high). Conversely, PtdIns(3,4,5)P_3_, which recruits, unfolds and activates Myo10 (*34, 53–55*), suits Myo10 perfectly because it represents a very small concentrated cellular compartment (*56, 57*). Recruitment and unfolding of Myo10 at PtdIns(3,4,5)P_3_ – rich leading edge and subsequent recruitment and concentration of Lpd at the leading edge (*35, 58*), promotes clustering of Ena/VASP proteins (*35, 44, 47, 48, 59*). Clustered VASP accelerates filament elongation while protecting nascent barbed ends from capping proteins, acting cooperatively with the WAVE regulatory complex (WRC) and Arp2/3 to drive localized protrusion (*38, 43, 49*). We propose that competing protrusions engage in a “winner-take-all” competition for a limiting pool of Myo10. Protrusions encountering permissive geometries recruit higher levels of Myo10, triggering a feed-forward amplification of actin assembly that stabilizes that front as the winner. In contrast, losing fronts fail to accumulate Myo10, stall, and trigger microtubule- and GEF-H1-dependent actomyosin retraction (*18, 21*). Without Myo10, no protrusion establishes dominance, thus no losing protrusion is established. Without that, the triggering of microtubule disassembly by the barrier effect, and the GEF-H1-dependent protrusion retraction pathway disassembly stimulates, will not occur. Consistent with this model, DCs depleted of Lpd, phenocopy Myo10 loss and exhibit defects in protrusion coordination.

Role for Ena/VASP proteins in this model is particularly attractive because their contribution to DC migration has been demonstrated directly (*60*). Combined Evl/VASP deficiency impairs several aspects – association with WRC components in DCs, and VASP KO DCS are sensitive to Arp2/3 inhibition, further suggesting functional cooperation between Ena/VASP-mediated filament elongated and Arp2/3 dependent actin assembly. Together with our Lpd depletion phenotype, these observations suggest that the Myo10-Lpd axis could spatially organize Ena/VASP and WRC/Arp2/3-dependent actin assembly at the dominant leading edge.

In summary, our findings establish that Myo10 functions as a spatial coordinator of dendritic cell migration. By defining the Myo10-Lpd-VASP axis as a critical governor of migratory persistence in complex environments, our studies links nanoscale cytoskeletal coordination to the macroscale efficiency of immune surveillance.

## Materials and Methods

### Reagents, Cell Culture and Molecular Biology

Primers were synthesized by Eurofins Genomics. Restriction enzymes, polymerases, cloning assembly kits and competent cells were from New England Biolabs and Thermo Fisher Scientific. Recombinant murine cytokines and chemokines were from Peprotech. HEK293T/17 cells were obtained from the American Type Culture Collection (ATCC) and authenticated by the vendor. Cells were maintained at 37°C in a 5% CO_2_ humidified incubator in DMEM supplemented with 10% FBS and 1% penicillin-streptomycin-amphotericin B (Thermo Fisher) and routinely checked for mycoplasma by PCR (Sigma-Aldrich). A complete list of reagents, chemicals, kits, antibodies and dyes is provided in Table S1.

All experiments involving recombinant DNA were conducted in accordance with the NIH guidelines for research involving recombinant or synthetic nucleic acid molecules. To generate lentiviral constructs of fluorescently-tagged tubulin and F-actin reporters, GFP-Tubulin and F-tractin mStayGold inserts were amplified by PCR and subcloned into BamHI/ EcoRI linearized pLV-IRES-blast vector using HiFi DNA Assembly (New England Biolabs). Clones were verified by colony PCR and diagnostic restriction digests, and resulting plasmids were verified by third-generation long-read sequencing (Plasmidsaurus).

Viruses were produced as previously described (*61*). Briefly, for production of retroviruses, transfer plasmids were co-transfected into 80% confluent HEK293T/17 cells with pCL-Eco helper plasmid at 1:1 ratio. Supernatants were harvested at 24 and 48 hours post-transfection, pooled, and filtered through a 0.45 µm PVDF filter. For production of lentiviruses, transfer plasmids were co-transfected into 80% confluent HEK293T/17 cells with psPAX2 and pMD2.G helper plasmids at 4.5:4.5:1 ratio. Supernatants were harvested at 48 and 96 hours post-transfection, pooled, filtered through a 0.45 µm PVDF filter, and concentrated using Amicon Ultra centrifugal filter units (100 kDa cut-off; Millipore Sigma).

For stable production of recombinant murine Flt3 ligand (Flt3L), a lentiviral transfer vector encoding secretion sequence of mouse Flt3L coupled to mCherry-P2A-puro sorting and selection cassette was synthesized by VectorBuilder, Inc. HEK293T/17 cells were transduced with the Ftl3L lentiviral supernatant, selected with puromycin and sorted for mCherry expression on BD FACSMelody sorter. Secreted Ftl3L concentration the conditioned media was quantitated using rat/mouse Flt3L ELISA kit (R&D).

### Animals

Animal protocols used in this study (H-0337 and H-2069) were reviewed and approved by the NHLBI Animal Care and Use Committee (ACUC). Animals were maintained at specific-pathogen-free animal facility in a 12-h light/ 12-h dark cycle with *ad libitum* access to water and food. Constitutive C57BL/6 *Myo10^-/-^* mice have been described previously (*62*).

C57BL/6 *Itgax-cre, -EFGP* (CD11c-Cre-GFP)(*63*) transgenic mice were obtained from The Jackson Laboratory. Wild type C57BL/6 controls were maintained in-house or purchased from Charles River laboratories. GFP-NM2A knock-in (GFP-NMHC II-A) (*64*) mice were backcrossed to pure C57BL/6 background for at least 10 generations. C57BL/6 *TcraTcrb* (OT-II) transgenic mice were obtained from the Jackson Laboratory as needed. Experiments were performed on age-matched and sex-matched 6- to 10-week-old male or female mice; littermate controls were used when possible. *In vivo* experiments were planned and executed to coincide with natural circadian rhythms of DC migration (Zeitgeber Time 7, or 7 hours after light onset) (*65*).

### Primary Dendritic Cell Isolation, BMDC Differentiation, and Transduction

For splenic DC isolation, spleens of WT and Myo10 KO mice were minced and dissociated in spleen dissociation medium (StemCell). DCs were isolated using negative selection pan-DC Enrichment Kit II (StemCell). For the isolation of dermal DCs, ears of euthanized mice were treated with depilatory cream (Nair), rinsed in PBS, split in dorsal and ventral halves, and incubated with Dispase II (1 U/mL; StemCell) for 45 min at 37 °C (*66*). Epidermal sheets were gently peeled away from the dermal sheets and discarded. Dermal sheets were further digested with Liberase TL (0.25 mg/mL; Roche) and DNAse I (0.2 mg/mL; Roche) for 90 min at 37 °C. Dermal suspension were filtered through 70-µm cell strainers, and total dermal DCs were enriched using CD11c Positive Selection Kit (StemCell). Isolated primary DCs were cultured overnight in complete R10 medium (RPMI 1640 + GlutaMAX, supplemented with 10% heat inactivated FBS, 10 mM HEPES, 1% penicillin-streptomycin, 1% amphotericin B, and 50 µM β-mercaptoethanol) containing 200 ng/mL LPS and 100 ng/mL Flt3L prior to functional assays.

Bone marrow-derived dendritic cells (BMDCs) were generated as described (*67*). Mononuclear cells were isolated from crude bone marrow extracted from tibiae and femurs of wild type and *Myo10^-/-^* mice and plated on non-tissue culture-treated dishes in complete R10 medium supplemented with 20% conditioned medium from a GM-CSF-secreting hybridoma (*68*) or 50 ng/mL recombinant GM-CSF. Fresh medium was added on day 3, 6 and 8 of culture. On day 10 the floating cell fraction was harvested gently without disturbing the attached cells and reseeded on tissue culture-treated dishes in complete medium supplemented with 200 ng/mL LPS. Floating cells were harvested the next day and used for flow cytometry analysis and experiments within 36 hours. Cells were routinely analyzed for the expression of CD11c, MHC-II, CD80, CD86 and CCR7 by flow cytometry. For transduction with retro- and lentiviruses, murine bone marrow mononuclear cells were transduced as described previously (*69*). Briefly, cells were transduced with retro- and lentiviruses plated on RetroNectin-coated plates at 900 x g for 45 min at 30°C. Cells were allowed to rest in the incubator for 2 hours before media change. Transduction was repeated 24-36 hours later. Cells were either selected with puromycin or FACS sorted.

### Flow Cytometry

Single cell suspensions were prepared in FACS buffer (2% heat-inactivated FBS, 25 µg/mL DNAse I, 0.5 mM EDTA in PBS) at 1 x 10^6^ cells/100 µL, blocked with anti-mouse CD16/32 and stained with fluorescently-conjugated antibodies on ice. For flow cytometric analysis of CCR7 expression, cells were stained at 37°C for 30 min with anti-CCR7 antibody prior to staining with other surface markers on ice. Cells were analyzed on Cytek Aurora spectral flow cytometer equipped with SpectroFlo software (Cytek Biosciences). Cells were gated according to forward- and side-scatter to yield singlets, followed by gating on viability dye-excluded cells, and further characterized by expression of surface markers. Data were analyzed with FlowJo (BD Biosciences).

### Contact Hypersensitivity and FITC painting

For the analysis of FITC-induced contact hypersensitivity, mice were sensitized topically on flanks with 50 µL of 1% (w/v) FITC dissolved in a 1:1 (v/v) mixture of acetone and dibutyl phthalate on days 0 and 6. On days 12, 13, and 14, sensitized mice were challenged on the dorsal side of an ear (one ear only) with 1% FITC. Ear thickness was measured using analog calipers.

For the analysis of *in vivo* DC migration using FITC painting, 20 µL of 1% (w/v) FITC in acetone and dibutylphthalate (1:1 ratio) was applied topically on the dorsal side of the ear of mice. Skin-draining auricular lymph nodes were harvested 48 hours later, digested in RPMI containing 0.05 mg/mL Liberase TL and 0.2 mg/mL DNAse I at 37 °C for 20 min, filtered through 40 µm strainer, counted and processed for flow cytometric analysis. Absolute counts were determined using CountBright counting beads (Thermo Fisher).

For competitive adoptive transfer studies, BMDCs generated from WT or Myo10 KO mice were pulsed with 200 ng/mL LPS for 2 hours, extensively washed, labeled with reciprocal combinations of dyes, and mixed at 1:1 ratio. A total of 1 x 10^6^ cells in a 25 µL volume of 0.9% (w/v) sterile saline were injected s.c. into a lateral tarsal region of the hind foot (one injection per recipient). Sham control recipients were injected with 25 µL saline only. Popliteal lymph nodes were harvested 24-48 hours later and either processed for flow cytometry or fixed and processed for cryosectioning and immunostaining. To assess the frequency of each BMDC genotype in the dLN via flow cytometric analysis, lymph nodes were teased through a 40 µm strainer and single cell suspension was stained with a viability dye, followed by CD45, CD11c, and MHC-II. Migratory DCs were identified as the CD11c^+^ MHC-II^hi^ population. Percentage of WT and Myo10 KO BMDCs in the dLN was calculated as a frequency of total migratory DC population. To assess the frequency by immunofluorescence staining, fixed dLNs were processed into 10 – 20 µm sections using Leica CM1860 UV cryostat and adhered to charged microscopy slides. Sections were permeabilized and blocked in 5% goat serum and 0.2% Triton-X100 in PBS and stained with an anti-laminin antibody overnight at 4°C in a humidified chamber. The next day, sections were washed thrice in 2% goat serum and 0.1% Triton-X100 in PBS and stained with a fluorescently-conjugated secondary antibody for 1 hr at room temperature. Post-incubation, sectioned were washed in PBS, mounted and imaged on Zeiss LSM 880 inverted confocal microscope using 20x/0.8 Plan-Apochromat objective. Large fields of view (800 µm x 800 µm) were acquired using tiling method (10% overlap) in the Zen software, followed by stitching.

### Ex vivo crawl-out assay

For determination of directed egress of DCs from the ear dermis, ears of euthanized mice were split into dorsal and ventral halves and floated dermis side down in 24-well plates in complete R10 medium supplemented with 600 ng/mL CCL21. After 48 hours, crawled out cells were pooled from the medium and stained for analysis by flow cytometry. Ears were digested in RPMI containing 0.25 mg/mL Liberase TL and 0.2 mg/mL DNAse I at 37 °C for 90 min, as described (*70*). Digested tissue was homogenized using manual tissue homogenizer, filtered through a 70 µm strainer and processed for flow cytometry staining and analysis. Absolute counts were determined using CountBright beads and the crawl-out efficiency was plotted as the ratio of emigrated or retained DCs relative to the total DC pool. For competitive crawl-in assays, ears from 6 – 8 week old mice were split into dorsal and ventral halves. Dorsal halves were pre-stained with LYVE-1 to mark lymphatics. BMDCs derived from WT and Myo10 KO mice were differentially labeled, mixed 1:1 ratio of 200,000 cells each and placed on top of exposed dermis for 15 min. Unbound BMDCs were rinsed off in R10 medium and ears were submerged into R10 medium and incubated at 37 °C for up to 3 hours. Ears were subsequently fixed in 2% PFA overnight at 4 °C, washed in PBS and mounted. Mounted specimens were imaged using confocal microscopy.

### Analysis of antigen uptake

For the assessment of gelatin degradation, immature (day 7) BMDC (200,000 cells) were plated on 0.5% glutaraldehyde-crosslinked Oregon Green - conjugated gelatin (0.2 mg/mL) in complete R10 medium supplemented with 50 ng/mL GM-CSF. Cells were allowed to degrade gelatin for 16 hours in a 37 °C tissue culture incubator. Post-incubation, chambers were washed with PBS and cells were fixed with 4% (w/v) PFA in PBS. Chambers were washed thrice in PBS and stained with Hoechst 33342 to mark nuclei and fluorescent phalloidin to mark filamentous actin. Chambers were scanned on Zeiss LSM 880 inverted confocal microscope via a tiling method (∼1000 µm x 1500 µm total area) using 20X objective. Endocytic capacity was evaluated by incubating immature DCs with Alexa Fluor 647– conjugated dextran or Alexa Fluor-conjugated ovalbumin (Thermo Fisher) at 37 °C or on ice (background) for 15 to 60 min, followed by cold washing and flow cytometric quantification.

### Microchannel Device Microfabrication

The master mold with 3.8 µm feature heights was created using standard photolithographic techniques (*71*). First, a 4-inch silicon wafer was dehydrated at 200 °C for 20 minutes. To pattern the features, SU-8 2005 was spin-coated onto the wafer. The wafer was then baked at 95°C for 2 minutes, patterned using a glass-chrome mask (Front Range Photomask) with a contact mask aligner (OAI Instruments) and LED illumination (Idonus). Next, the wafer underwent a post-exposure bake at 95 °C for 2.5 minutes, after which it was developed in a SU-8 developer bath with gentle agitation for approximately one minute, before a rinse in isopropanol, water, and then drying. To enhance durability, the master mold was hard baked at 200 °C for 4 minutes and then cooled down to room temperature at a rate of 150 °C per hour. Finally, the wafer was silanized overnight with tridecafluoro-1,1,2,2-tetrahydrooctyl-1-trichlorosilane vapor to assist with the unmolding process. PDMS was prepared by mixing the base and curing agent in a 10:1 weight-to-weight ratio using a planetary mixer (Thinky). The mixed PDMS was then poured onto the master mold, degassed to remove air bubbles, and cured at 80 °C for one hour. After curing, the PDMS was carefully removed from the mold. Inlet and outlet reservoirs were created using a 2 mm biopsy punch (Acuderm), and the individual devices were cut out using an 18 mm arch punch (Osborn). On the day of the experiment, microfluidic devices and the 35 mm glass-bottom dishes were plasma treated with oxygen plasma for 35 seconds using the plasma cleaner (Thierry Plasma) and immediately assembled by gently pressing the PDMS device onto the dish. To improve bonding, the assembled devices were incubated at 80 °C for 5 minutes before filling the channels with 5% BSA in PBS supplemented with 1x penicillin-streptomycin and 1% amphotericin B to maintain the hydrophilicity.

### Under agarose migration assay

The under agarose migration assay was adapted from (*72*). Briefly, 4-well glass-bottom chambers (CellVis) were blocked with 5% BSA or 0.1 M PLL-PEG overnight. Agarose layer was prepared by mixing 2% UltraPure Agarose (Invitrogen) melted in HBSS and complete R10 medium supplemented with 20% FBS in a 1:1 ratio, resulting in 1% agarose in complete R10 medium with 10% FBS. Inlet and outlet reservoirs were created using a 1 mm biopsy punch (WPI) and were spaced 2 mm apart. CCL19 (600 ng/mL) in phenol red-free R10 medium was added to the first port and the chamber was incubated for 30 min at 37 °C before loading cells (50,000 cells) into the second port. The chamber was incubated at 37 °C for 2-3 hours prior to live imaging or fixation for immunofluorescence staining.

### Migration in 3D matrices

Collagen gel mixture was prepared as described (*67*). Briefly, rat type I collagen (Corning) was mixed on ice in 10X MEM and 7.5% sodium bicarbonate taking care of achieving a final pH of 7.3-7.5. Gel mixture was immediately mixed with cells resuspended in complete R10 medium for a final collagen concentration of 1.6 mg/mL. The mixture was loaded into a custom glass-bottom chamber. Following polymerization at 37 °C for 45 min, CCL19 (600 ng/mL) was overlaid to create a gradient. For collagen invasion assays, cell-free collagen gels containing 600 ng/mL of CCL21 were cast on glass-bottom chambers.

After polymerization, the gels were overlaid with 100,000 – 200,000 BMDCs in complete medium and incubated overnight in tissue culture incubator. Gels were fixed with 4% PFA and imaged using confocal microscopy to determine invasion depth.

### Immunofluorescence staining

Cells were fixed rapidly using pre-warmed Zamboni fixative (2% (v/v) methanol-free PFA, 0.2% picric acid in PBS) or pre-warmed 4% (v/v) methanol-free PFA in PHEM/sucrose buffer (10 mM PIPES, 10 mM HEPES, 100 mM NaCl, 1 mM EGTA, 3 mM MgCl_2_, and 0.3 M sucrose) at 37 °C for 10 min, followed by a 10-min fixation at room temperature. Chambers were then washed thrice in PBS, permeabilized with 0.1% Triton-X100 in PBS for 5 min, and blocked with 5% (v/v) goat serum in PBS for 30 min. Chambers were stained overnight at 4 °C with antibodies diluted in 2% BSA in PBS. The next day, chambers were washed thrice in PBS and stained with fluorescent dye-conjugated secondary antibodies and/or phalloidins at room temperature for 1 h. The chambers were washed thrice in PBS and imaged by confocal or TIRF microscopy.

### Immunoblotting

Cells were lysed in lysis buffer supplemented with phosphatase and protease inhibitor cocktail (Cell Signaling) by sonication. Lysates were cleared by centrifugation at 10,000 x g for 5 min and quantitated for total protein content by the BCA assay (Pierce). Lysates were denatured at 95 °C for 3 min in Laemmli sample buffer supplemented with 355 mM 2-mercaptoethanol and resolved on discontinuous 6% Tris-Glycine SDS-PAGE (Bio-Rad) gels and transferred onto nitrocellulose membranes using Bio-Rad Turbo Transfer apparatus.

Membrane slices were blocked in 3% milk in TBS for 1 hr at room temperature and incubated in primary antibodies diluted in 5% milk in TBS-T (Tween-20 0.05%) overnight at 4°C. The next day, membrane slices were washed thrice in TBS-T and incubated with appropriate HRP-conjugated secondary antibodies diluted in 5% milk in TBS-T for 1 hr at room temperature.

Membrane slices were washed twice in TBS-T, once in TBS and developed using Clarity ECL reagent (Bio-Rad) on Amersham GE AI600 imager. Chemiluminescent signals were quantitated manually in Fiji by normalization to the loading control and subsequent comparison of the treatment sample to the control sample. For a complete list of reagents and antibodies used for SDS-PAGE and immunoblotting, please see Table S1.

### Microscopy

Live cell time-lapse imaging was performed on a Nikon A1R inverted confocal microscope equipped with an environmental stage incubation enclosure (Tokai Hit) using a Plan Fluor 20 x /0.75 NA objective in resonant scanning mode. Frames were recorded every 2 min for 3-4 hours. Confocal imaging was performed using Zeiss LSM 780 using a Plan Apochromat 40 x /1.4 NA oil objective. High-resolution imaging of under agarose migration was performed using Zeiss LSM 880 AiryScan using a Plan-Apochromat 60 x /1.49 NA oil-immersion objective or DeltaVision OMX system (GE Healthcare) using a 60x /1.42 NA objective (Olympus). Airyscan image reconstruction was performed using Zeiss ZEN imaging software.

### Image Analysis

Fiji (NIH), Excel (Microsoft), custom Python code, Chemotaxis Tool (Ibidi), and Prism (GraphPad Software) were used for image analysis. To quantitate cell migration parameters, cell nuclei were tracked using “Manual Tracking” plugin in Fiji. The resulting tracks were imported into Chemotaxis Tool (Ibidi) for further analysis. Custom computational pipelines for leading-edge stability and PtdIns(3,4,5)P_3_ temporal alignment analyses were developed with assistance from OpenAI HHS Enterprise ChatGPT (GPT-5.6 Sol) through iterative process of code debugging and workflow design by the authors.

Leading edge dynamics were quantitated from time-lapse CellMask plasma membrane stain movies using a custom Fiji macro. Individual cells were followed every frame and each frame was manually classified as containing one dominant front, two or three and more competing fronts. The location of the dominant front was simultaneously assigned to one of the eight 45° sectors defined relative to the cell centroid. The multifront fraction was calculated as the fraction of annotated frames containing two or more simultaneous fronts. Spatial leading-edge stability was quantitated as the normalized Shannon entropy of dominant-front occupancy across the eight sectors,

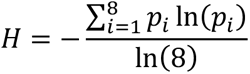

where *p_i_* is the fraction of observations in which the dominant front occupied sector *i*. Thus, values approaching 0 indicate confinement of the dominant front to a consistent cellular region, whereas increasing values indicate broader spatial redistribution of leading-edge activity. Temporal alignment analysis of PtdIns(3,4,5)P_3_ quantitated from TIRF time-lapse movies using a custom Python analysis pipeline. For each cell, a PtdIns(3,4,5)P_3_-enriched region was assigned manually in the first frame and subsequently tracked relative to the cell centroid. Cell-centroid trajectories were median-smoothed over a 1-s window, and movement direction was calculated from displacement over a 2-s interval. To determine whether PtdIns(3,4,5)P_3_ polarity predicted subsequent cell movement, the angular orientation of the PtdIns(3,4,5)P_3_-enriched region at time *t* was compared with the direction of movement beginning at time *t* + *τ*, where *τ* was varied from −10 to +10 s in 0.2-s increments. Directional alignment was calculated as the cosine of the angular difference between PtdIns(3,4,5)P_3_ polarity and movement direction,

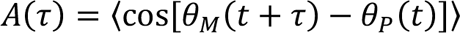

where *θ_P_* is the cell-centered orientation of PtdIns(3,4,5)P_3_ enrichment and *θ_M_* is the direction of cell movement. Values approaching 1 indicate movement aligned with PtdIns(3,4,5)P_3_ polarity, values near 0 indicate no directional relationship, and negative values indicate movement oriented opposite to PtdIns(3,4,5)P_3_ polarity. Positive lag values therefore indicate the extent to which PIP3 polarity predicts subsequent movement.

### Statistical Analysis

Statistical analysis was performed using Prism 10 (GraphPad). Comparison of two groups was carried out using Student’s t-test, comparison of datasets with more than two groups was carried out using One- or Two-way ANOVA, as appropriate, with recommended multiple comparisons tests. Animal studies were planned to provide 60-80% power for a target effect size of 1.2-1.5. See figure legends for specific sample sizes, denoted as “N.” Alpha was set to 0.05. Data are presented as mean ± s.e.m., unless otherwise noted in the legend. Results for statistical tests are presented as p-value nominations, where * corresponds to p ≤ 0.05, ** to p ≤ 0.01, *** to p ≤ 0.001, **** to p < 0.0001.

Migration experiments shown in Figures S3 and S4 were independently performed and analyzed by N.M.B. and A.G. without their prior knowledge of genotypes and conditions. In addition, N.M.B. performed secondary blinded scoring of leading edge organization shown in Figure 4.

## Data availability

Data are available in the article itself and its supplementary materials.

## Acknowledgements

We would like to thank NHLBI Animal Surgery and Resource for advice and training, NHLBI Flow Cytometry Core CRC section for training, NHLBI Light Microscopy Core deputy director Dr. Xufeng Wu for training, and NIAMS Light Microscopy Core for the use of Leica cryostat. We would like to additionally thank Drs. Yasmine Belkaid (Institute Pasteur), Jose Cancelas (Dana-Farber Cancer Institute), Nicolas N. Nassar (Cincinnati Children’s Hospital Medical Center), Paul A. Randazzo (National Cancer Institute), and Vanja Lazarevic (National Cancer Institute) for helpful discussions and for sharing technical expertise and reagents.

This work was funded by the Intramural Research Program of the National Heart Lung and Blood Institute of the National Institutes of Health to J.A.H., the NHLBI Lenfant Postdoctoral Fellowship to A.G., and NHLBI K22 Career Transition Award to A.G.

This research was supported by the Intramural Research Program of the National Institutes of Health (NIH). The contributions of the NIH authors are considered Works of the United States Government. The findings and conclusions presented in this paper are those of the authors and do not necessarily reflect the views of the NIH or the U.S. Department of Health and Human Services.

## Author contributions

Conceptualization – A.G. and J.A.H.; Methodology – A.G., P.R.S., N.Y.M., and J.A.H.; Validation – A.G., P.R.S., and N.M.B.; Formal analysis – A.G. and N.M.B.; Investigation – A.G.; Resources – A.G., P.R.S., Y.Y. and D.C.S.; Data curation – A.G.; Writing – original draft – A.G. and J.A.H.; Visualization – A.G.; Funding Acquisition – J.A.H.; Supervision – N.Y.M. and J.A.H.

## Disclosures

The authors declare no competing interests.

## Notes

### Competing Interest Statement

The authors have declared no competing interest.

