## Supplementary figures and images for "Myo10-dependent leading-edge coordination maintains physical coherence of migrating dendritic cells"

### Supplemental_Figures

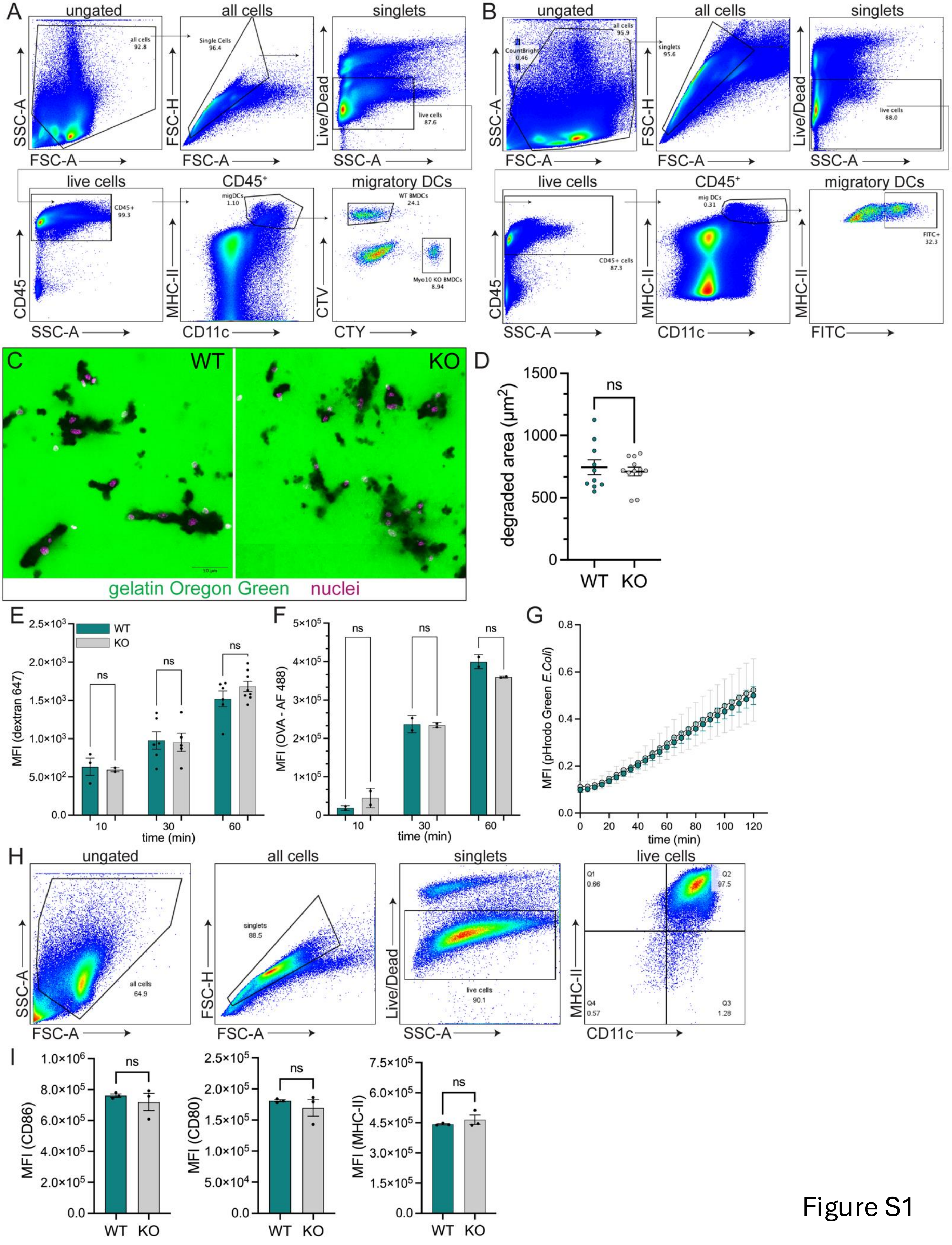

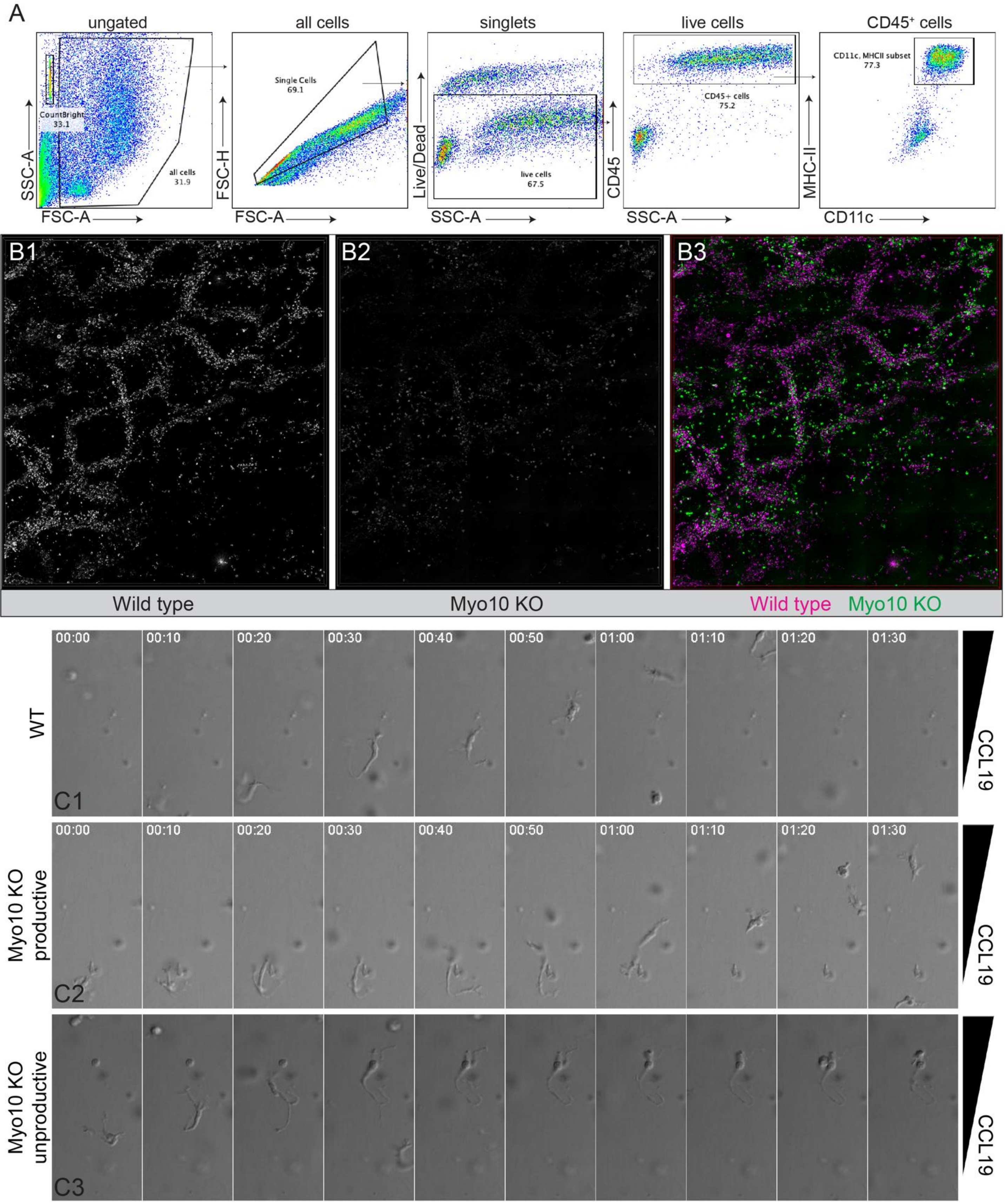

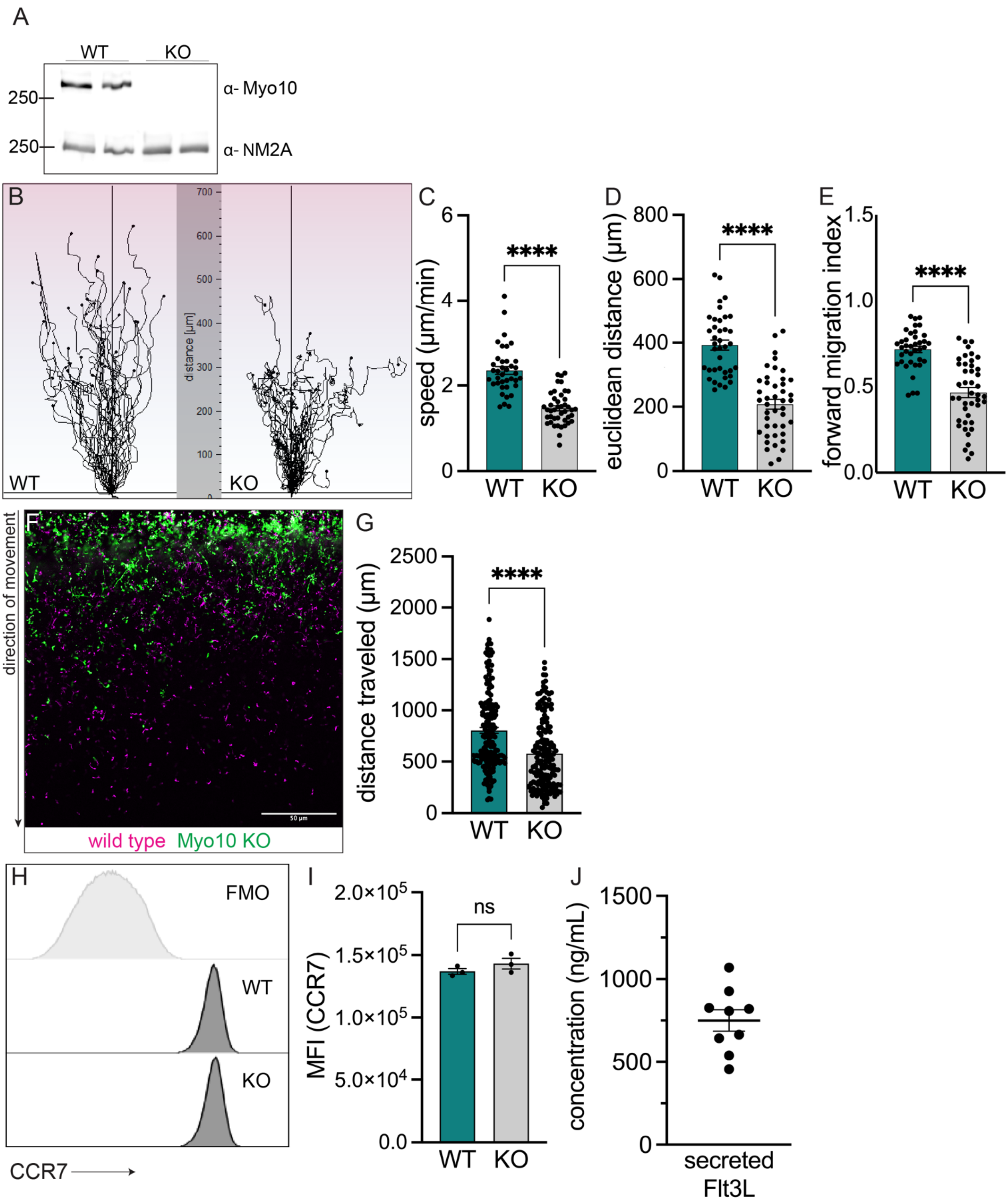

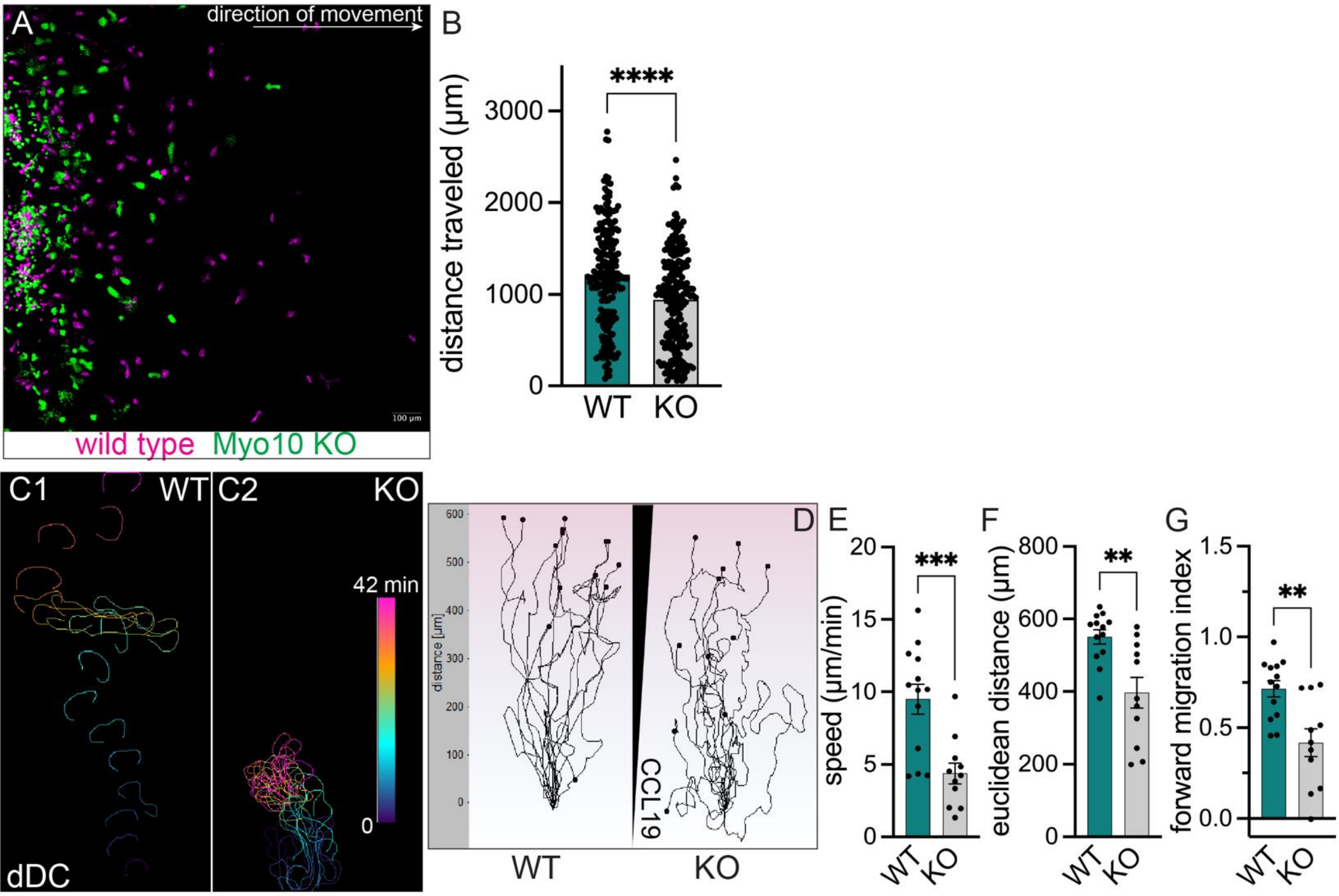
